# Transposable elements resistant to epigenetic resetting in the human germline are epigenetic hotspots for development and disease

**DOI:** 10.1101/2020.03.19.998930

**Authors:** Sabine Dietmann, Michael J Keogh, Walfred Tang, Erna Magnusdottir, Toshihiro Kobayashi, Patrick F Chinnery, M. Azim Surani

## Abstract

Despite the extensive erasure of DNA methylation in the early human germline, nearly eight percent of CpGs are resistant to the epigenetic resetting in the acutely hypomethylated primordial germ cells (week 7-9 hPGCs). Whether this occurs stochastically or represents relatively conserved layer of epigenetic information is unclear. Here we show that several predominantly hominoid-specific families of transposable elements (TEs) consistently resist DNA demethylation (henceforth called hPGC-methylated TEs or ‘escapees’) during the epigenetic resetting of hPGCs. Some of them undergo subsequent dynamic epigenetic changes during embryonic development. Our analysis of the fetal cerebral cortex also revealed multiple classes of young hPGC-methylated TEs within putative and established enhancers. Remarkably, specific hPGC-methylated TE subfamilies were associated with a multitude of adaptive human traits, including hair color and intelligence, and diseases including schizophrenia and Alzheimer’s disease. We postulate that hPGC-methylated TEs represent potentially heritable information within the germline with a role in human development and evolution.

## INTRODUCTION

The unique and comprehensive erasure of DNA methylation in early human primordial germ cells (hPGCs) during their migration into the developing gonads is a prerequisite for totipotency of zygotes, through which the germline transmits both genetic and epigenetic information for the development of subsequent generations. Recently, however, we identified TE subfamilies consisting primarily of hominoid-specific TEs, which are resistant to DNA demethylation during the epigenetic resetting leading to acute hypomethylation of hPGCs at ~ week 7-9 of development^1^.

Approximately forty percent of the human genome consists of TEs^2,3^. Whilst older TEs have lost the ability to mobilise within the genome, young TEs can potentially retrotranspose, especially in the germline and during early development, when extensive epigenetic reprogramming occurs. To counter their activity, host defense mechanisms, including DNA methylation and piRNA biosynthesis, ensure their repression. The acute loss of DNA methylation in PGCs precedes the onset of piRNAs biosynthesis that could therefore potentially lead to the activation of TEs. Another mechanism for host defense might involve Krüppel-associated box (KRAB) zinc finger proteins (KZFPs) to suppress TEs in hPGCs. KRAB-ZFPs can bind to genomic retrotransposons, and transcriptionally silence them via the recruitment of repressive complexes through KAP1^4^. In other contexts, such as during embryonic development, KZFP-TE interactions might influence the expression of surrounding genes^4–6^, and play a role in regulating development. Accordingly, there is a significant potential for the involvement of this large and rapidly evolving family of KZFPs during development^7^.

Approximately ninety-five percent of TEs have evolved neutrally from their ancestral origin^8^, though some intergenic TEs appear to be under purifying selection for a functional role^9^; for example, those located close to genes involved in human development^10^. Such positively selected TEs have undergone exaptation and acquire functions such as cis-regulatory elements (CREs)^11, 12^, and recruit RNA polymerases to act as promotors^13^. Mutations within TEs can generate novel transcription factor (TF) binding sites^14^, and conversely, mutations in TFs can alter their functions. The acquisition of mutations within TFs and their binding sites, such as in TEs, might promote novel gene regulatory networks for for mammalian development^15^. Recent studies suggest that the interaction between KZFPs and TEs that have arisen within our hominoid ancestry might facilitate their development as *cis* regulators affecting early and late embryogenesis^16^. Accordingly, KZFP-TE mediated chromatin modifications have the potential to affect nearby genes within the developing fetus in tissue and cell-specific manner; this may account for species-specific epigenetic and transcriptional networks even when they share broadly similar genomes^16^,^17–19^. What drives or promotes the formation of new gene regulatory networks remains to be fully elucidated.

Here we focus on the DNA demethylation-resistant hominoid-specific TEs in hPGCs and ask whether their presence within the germline might have contribute to human development. By considering the age at which incorporation of each TE occured into the human genome^19^ and their putative biological effects, we explore how they might have facilitated specific facets of human development. In particular, we have considered this possibility in the context of human neurological development.

## RESULTS

### The spectrum of transposable elements resistant to DNA methylation erasure in hPGCs

First, we examined the genomic regions that are resistant to DNA methylation erasure within hPGCs (in male and female week 7-9 hPGCs), when they reach an unprecedented acute hypomethylated state^1^ (Fig. 1a). Using a hidden Markov model (HMM), we identified 116,617 significantly hypermethylated genomic regions, with average methylation of > 30%; 60,657 of them overlap with those from an independent study of week 10-11 hPGCs^21^ (Fig. 1b, Extended Data Fig.1a, Supplementary Table 1).

**Fig. 1:**
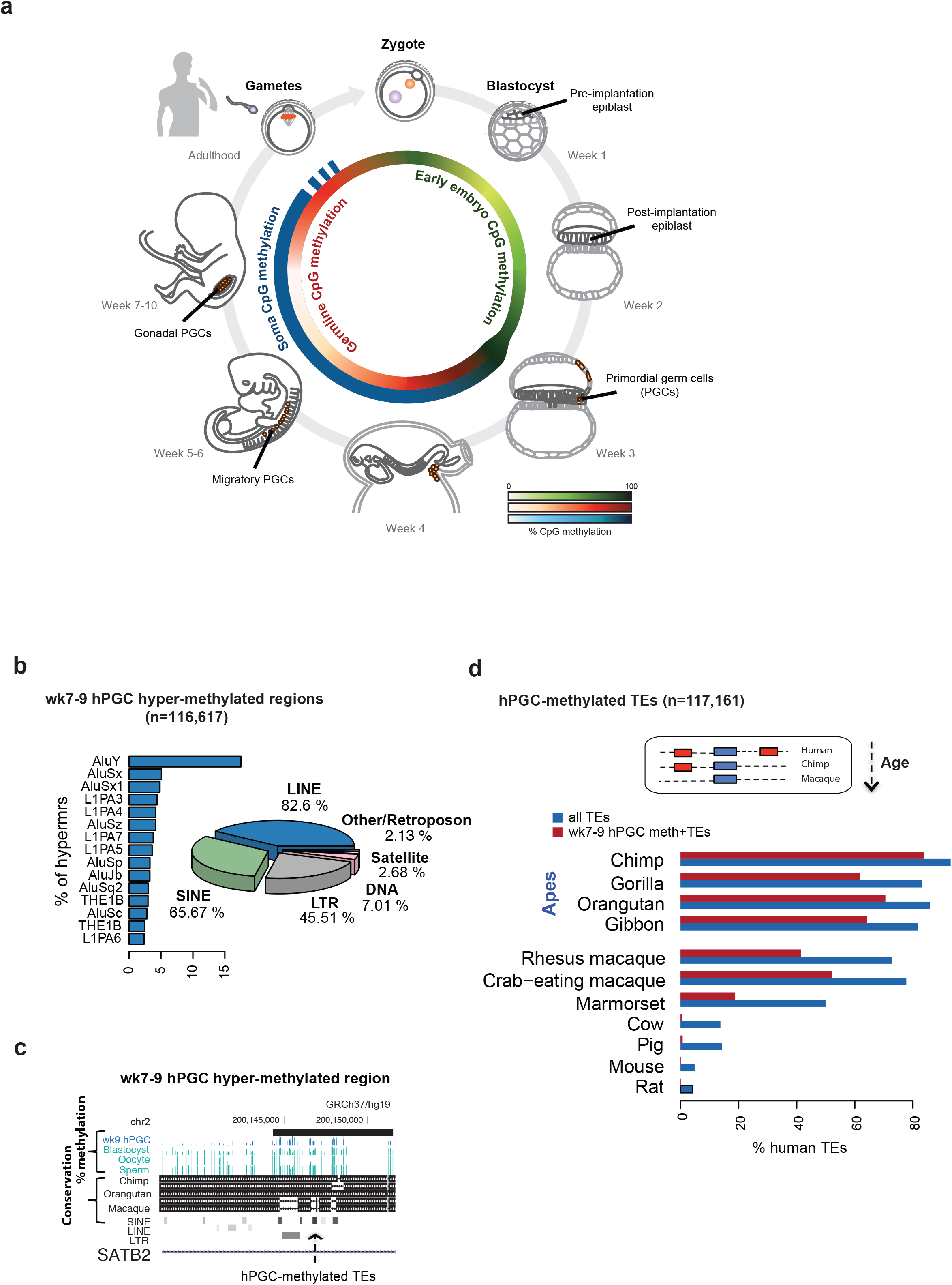
The potential of TE elements for transgenerational inheritance. **a,** A schematic representing the stages of the human germline cycle. The inner circle indicates the average % DNA methylation level of TE elements at those stages. The outer circle indicates the % DNA methylation in somatic tissues. **b,** Pie chart and bar plot showing the percentage of elements, which intersect with 116,617 significantly hyper-methylated regions in wk7-9 hPGCs for each TE subfamily indicated. TE elements were defined as those that cover at least 10% of the length of the hyper-methylated region (see also Supplementary Table 1). **c,** A schematic illustrating a hyper-methylated region (“escapee”) within the intron of the SATB2 gene, which overlaps with multiple TE elements. The region’s % DNA methylation in wk9 hPGCs and gametes, and its conservation in non-human primates is shown. **d,** A bar plot showing the percentage of TE elements in the human genome that are conserved in non-human primate and mammalian genomes (defined as TEs where at least 80% of the human TE element can be mapped to a syntenic region in the other species by liftOver). For each species, bars represent TE elements that are methylated in wk7-9 hPGCs (DNA methylation > 30%) in red, and all annotated TE elements in blue (see also Supplementary Table 2). Whilst 75-80% of non-methylated human TE elements are detected in apes and macaque, hPGC-methylated TE elements are more preferentially conserved in apes.

We detected 117,161 transposable elements within the hypermethylated genomic regions (subsequently termed ‘hPGC-methylated TEs’, Fig. 1c), many of which were hominoid-specific (< 25 million years old) and shared with apes. While a lower proportion of them was also present in macaques, a negligible proportion was found in distant mammals as compared to all TEs (Fig. 1d, Extended Data Fig. 1c, Supplementary Table 2). A smaller number of older TE families of >25 million years, however, also remained methylated (Extended Data Fig. 1b,c; Supplementary Table 2,3).

The hPGC-methylation of TEs varies markedly by TE family, with Alu (AluY, AluSx; members of the SINE family), long-terminal repeat (LTR) THE1B, together with the long interspersed nuclear element (LINE) L1PA3-7 subfamilies of retrotransposons (Fig. 1b) being particularly abundant within hypermethylated regions. The proportion of TEs within these subfamilies that are methylated also appear to be proportional to the age of the TE subfamily. For example, in the hominid-specific subset of retrotransposons, SINE-VNTR-Alu (SVA), SVA_A (the oldest SVA originating ~15 million years ago in early primate evolution^22^) had a far higher percentage of methylated elements (~ 80%), compared to the much younger human-specific SVA_E (3.5 million years old^22^), with only ~ 25% of methylated elements (Extended Data Fig. 1b,c). Similarly, in LINE1 family, where L1PA3-7 are between 12.5 and 31 million years old, older TEs (e.g. L1PA7) show a greater proportion of TEs remaining methylated compared to younger TEs (e.g. L1PA3). This is observed even though all these TE subfamilies are inactive in humans^22^. Other very young TEs, such as multiple members of the AluY family, which arose < 3 million years ago within humans^24^, remain largely unmethylated even though they have the potential to mobilize (Extended Data Fig. 1b,c).

Why a progressive increase in DNA methylation occurs with the age of TEs, and whether it has any functional significance is unknown. However, it suggests that the presence of hPGC-methylation of TEs may not primarily function to reduce TE activity, as transposition activity decreases with TE age. Since the acute DNA demethylation of week 7-9 hPGCs occurs before the onset of Piwi-RNAs (piRNA)^25–27^, (another key host defense mechanism), implicates other mechanisms, such as histone modifications to counter potential TE activity. In this regard, KZFP-TE interactions also merit consideration for regulating potentially active and very young TEs (< 1 million years old) during the hypomethylation of week 7-9 hPGCs (see below).

### Tracing the progression of hPGC-methylated TEs at the onset of development

To begin to determine whether hPGC-methylation of TEs may therefore function to promote histone modifcations within human tissues (rather than repress TE activity within the germline), we determined the overlap between hPGC methylated TEs and multiple histone modifications present within 37 primary tissues representative of all significant human lineages^28^. We observed that hPGC-methylated TEs intersected H3K9me3 marks significantly more frequently (58%) than non-methylated TEs (25%) in the genome (P value < 2e-16, Pearson’s Chi-squared test) (Fig. 2a).The enrichment of H3K9me3 marks in hPGC-methylated TEs appears to arise at different rates between different TE subfamilies. For example, hPGC-methylated SVAs are enriched for fetal H3K9me3 marks within ~3.2 million years (SVA_F), whereas the youngest LINE1 hPGC-methylated TEs acquire H3K9me3 marks within ~ 7.6 million years^29^ (Fig. 2b, Extended Data Fig. 2a). Importantly, these timeframes also reflect the rate at which DNA methylation is acquired within hPGCs by these families (Fig. 2c). We also observe an enrichment of H3K9me3 on human-specific hPGC-methylated TEs in the fetal and adult brain (see below, Fig. 2b, Extended Data Fig. 2a). These data further support the hypothesis that hPGC-methylated TEs may mark or even influence the acquisition of H3K9me3 in human tissues. Moreover, some of the TE subfamilies with the highest fraction of hPGC-methylated elements, including LTR12, MER11A-B. LTR12C, and the SVAs, were significantly enriched for tissue-specific enhancer marks (K4me1+K27ac) (P value < 1e-5, Pearson’s Chi-squared test, Extended Data Fig. 2b, Supplementary Table 4).

**Fig. 2:**
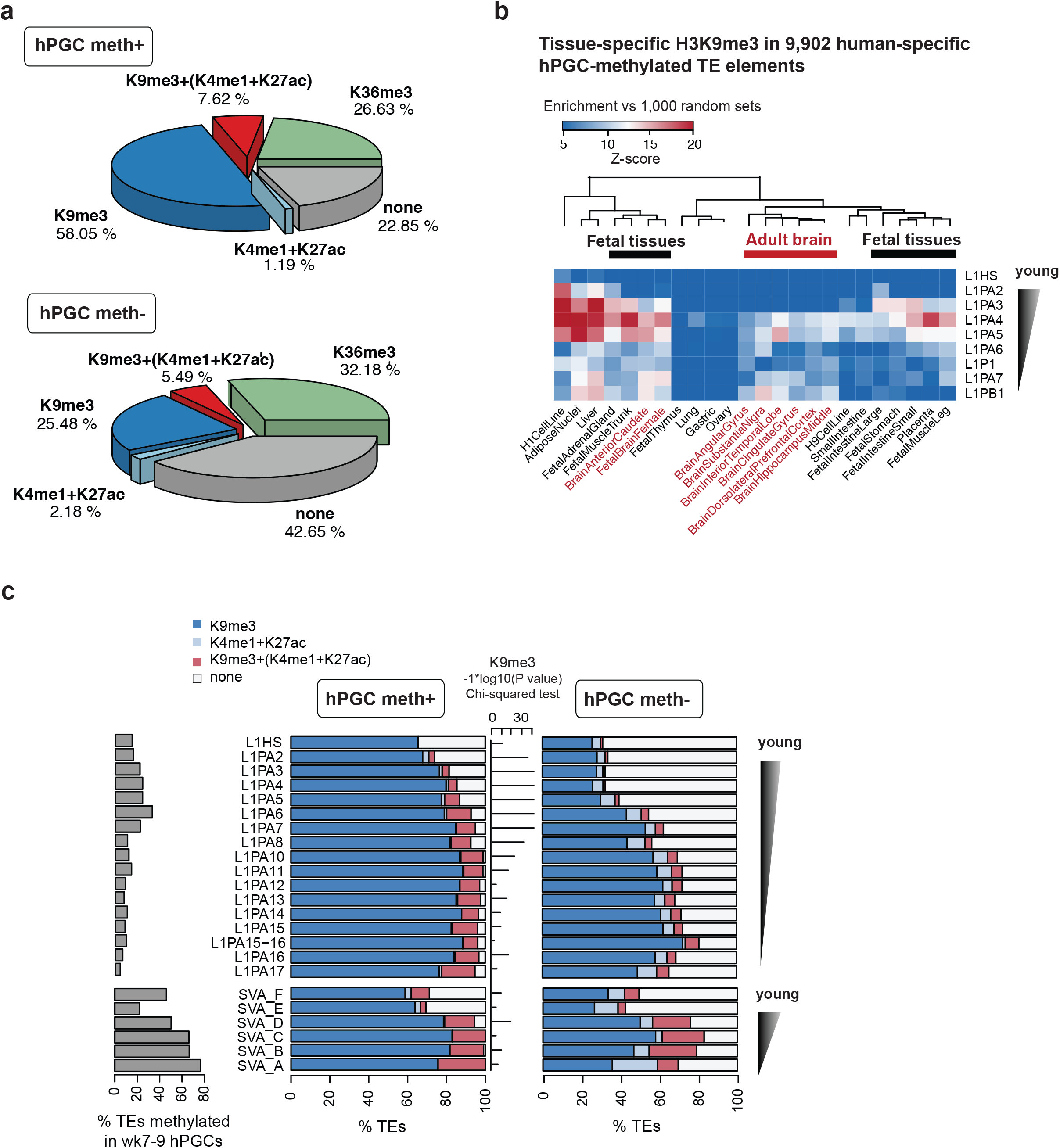
Association between hPGC-methylated regions and tissue-specific histone marks. **a,** Pie charts showing the percentage of human TE elements that intersect with established histone modification peaks in 7 fetal and 28 adult tissues (data from NIH Roadmap Epigenomics Project). The percentage of TE elements are shown for (top) those that are methylated in wk7-9 hPGCs (% DNA methylation > 30%) and (bottom) those that are non-methylated. **b,** A heat map depicting the enrichment (Z-scores) for tissue-specific H3K9me3 ChIP-seq peaks (data from NIH Roadmap Epigenomics Project) in 9,902 human-specific hPGC-methylated TE elements, compared to 1,000 random sets. Shown are the human-specific TE elements that cannot be mapped to syntenic regions in non-human primates or other mammals, and with an enrichment Z-score of at least 10 for one cell type. **c,** Bar plots showing the percentage of TEs in each subfamily that intersect with H3K9me3 and enhancer-related (H3K4me1+H3K27ac) ChIP-seq peaks in 7 fetal and 28 adult tissues (data from NIH Roadmap Epigenetics Project, see also Supplementary Table 4). TE elements are categorized as methylated (> 30 % DNA methylation) or non-methylated in wk7-9 hPGCs. LINE L1 and SVA subfamilies are ranked according to their age. Log10-transformed P values are indicated by bars. Younger primate-specific LINE L1 subfamilies (L1PA2-L1PA8) – but not the youngest (L1HS) - display the most significant differences in H3K9me3 for methylated vs non-methylated TEs. Left: bar plot showing the fraction of TE elements for each subfamily that are methylated in wk7-9 hPGCs (DNA methylation > 30%).

To try and gain insigt into how hPGC-methylation may influence the potential development of both H3K9me3 and enhancer histone modifications, we considered the fate of hPGC-methylated TEs during subsequent development. Beginning with gametogenesis, we observed that the majority of older hPGC-methylated TEs retain methylation during gametogenesis in a relatively homogeneous fashion. The youngest TEs however (including AluYa5, AluYg6, LTR12C-F, and SVA B-F), showed significant DNA methylation heterogeneity within oocytes, and more markedly within sperm (Extended Data Fig. 3a-c, Supplementary Table 2,5). Consequently, some young TE subfamilies may be a source of epigenetic heterogeneity in sperm, while older methylated TEs are likely to be conserved and consistently transmitted by sperm and oocytes.

Since it is technically challenging to follow the fate of hPGC-methylated regions during early human development, we considered a surrogate model of *in vitro* transition of human embryonic stem cells (hESCs) from naïve to primed, representing the preimplantation epiblast, and the more advanced pre-streak gastrulating postimplantation epiblast, respectively^30,–32^ (Fig. 3a). We observed overall a correlation of DNA methylation levels in hPGC-methylated TEs between week 7-9 hPGCs and the preimplantation epiblast (Extended Data Fig. 3b); the hPGC-methylated regions showed distinct dynamic changes concerning H3K9me3 and DNA methylation during the transition to primed hESCs. Evidently, some TEs such as SVA_D and SVA_F show marked increases in H3K9me3 levels in primed hESCs in both hPGC-methylated and non-methylated TEs. Conversely, we observed a loss of H3K9me3 on the hPGC-methylated elements of most TE subfamilies in primed hESCs, while there is a gain of H3K9me3 on unmethylated TEs (Fig. 3b,c, Extended Data Fig. 3d). A fraction (5-10%) of hPGC-methylated TEs that lose H3K9me3 in primed hESCs intersects with established tissue-specific enhancer signatures (Fig. 3b,c), including TE elements of the LTR1D, LTR12C, HERV9 and HERVH subfamilies^33,34^. In contrast, hPGC-methylated TEs that gain H3K9me3 during the naïve-to-primed transition generally lack tissue-specific enhancer marks (Fig. 3b).

**Fig 3:**
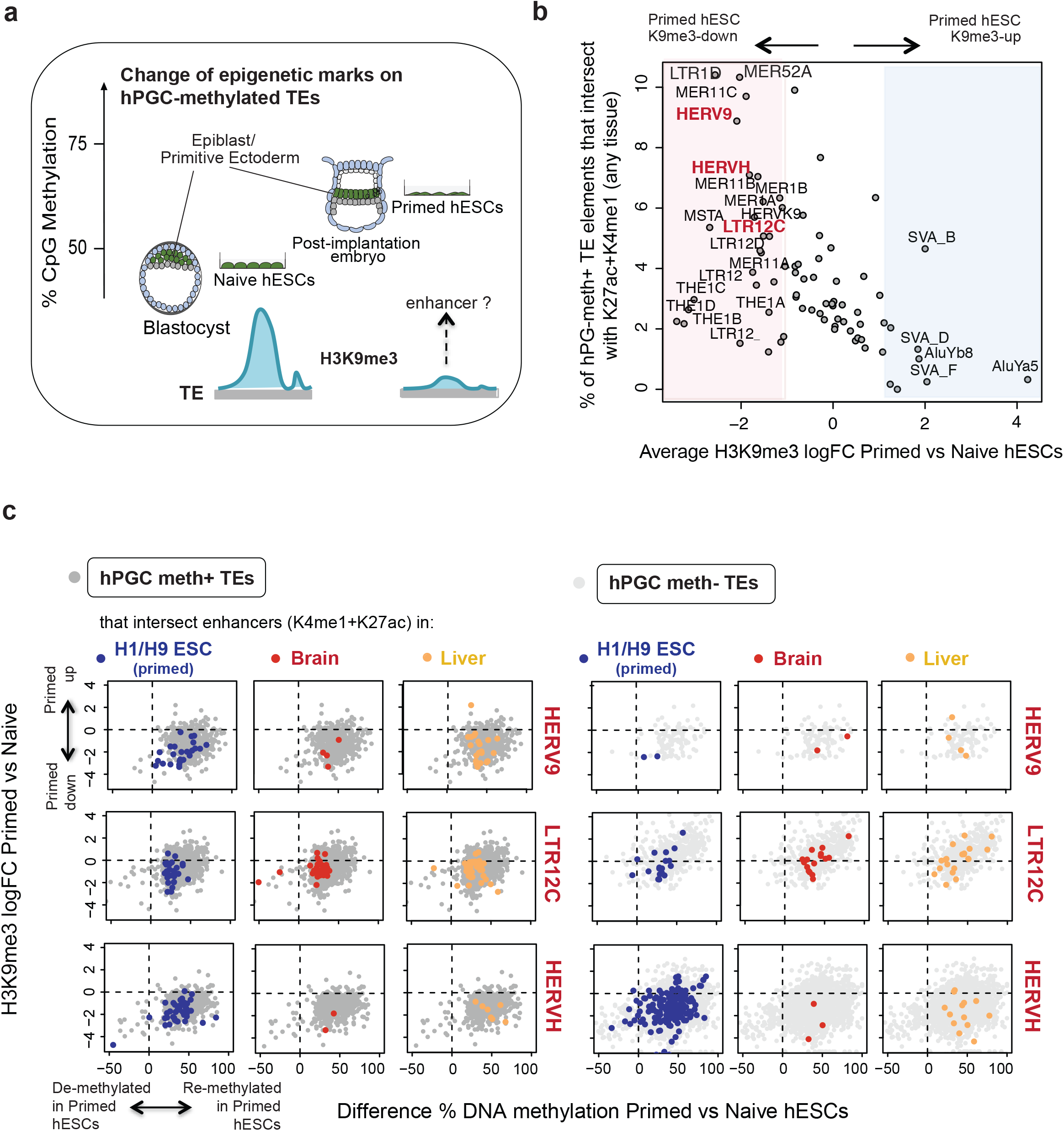
Dynamic of epigenetic marks on hPGC-methylated TE elements during early human embryogenesis. **a,** A schematic showing the % DNA methylation levels of *in vitro* hESCs and the most comparable human embryonic developmental stages. The analyses at these developmental stages characterize hPGC-methylated TE elements that lose H3K9me3 during the transition from naïve to primed hESCs. Some TE elements subsequently gain H3K27ac and can act as enhancers at later developmental stages. **b,** A scatter plot showing for each TE subfamily the percentage of hPGC-methylated elements that intersect with enhancer-related (H3K4me1+H3K27ac) ChIP-seq peaks in any of the 36 *in vivo* cell types (data from NIH Roadmap Epigenomics Project), compared to the average H3K9me3 log fold change (logFC) in primed vs naïve hESCs (data from Theunissen et al. 2016). The names of the TE subfamilies for which the DNA methylation and H3K9me3 dynamics is shown in Fig 3c are printed in red. With the exception of SVAs, which gain H3K9me3 on average, most TE subfamilies lose H3K9me3 in primed hESCs. 6-12% of the TE elements of characteristic subfamilies, including LTR1D, HERV9 and HERVH, intersect with active enhancer marks. **c,** Scatter plots of the H3K9me3 logFC compared to the difference in % DNA methylation in primed vs naïve hESCs (data from Theunissen et al. 2016, Guo et al., 2017) for TE elements that are methylated (> 30 % DNA methylation) or unmethylated in wk7-9 hPGCs. TEs that intersect with cell type-specific enhancers (H3K4me1+H3K27ac ChIP-seq peaks, data from NIH Roadmap Epigenomics Project) are shown in colour as indicated above the plots. Most TEs gain DNA methylation in primed hESCs while progressing from de-methylated naïve hESCs. In contrast, hPGC-methylated TE elements lose H3K9me3 and become enhancers more frequently than unmethylated TEs, while non-methylated TEs gain H3K9me3 (see also Extended Data Fig. 3d).

The loss of H3K9me3 enables lineage selection in early progenitor cells, accompanied by more dynamic changes than was thought previously^35^. Observations of early murine embryos, for example, have also shown that H3K9me3 may replace DNA methylation on TEs in a family-specific manner during preimplantation development^36^. H3K9me3 in the preimplantation epiblast, with its widespread DNA hypomethylation, might therefore ‘mark’ these TEs before DNA re-methylation occurs in the postimplantation epiblast. Irrespective of whether H3K9me3 is enriched or lost, methylation of TEs in hPGCs appears likely to influence the histone modification dynamics within TE families during naive to primed transition. Notably, a reduction in H3K9me3 levels on the majority of TE families is associated with lineage-specific enhancers.

Thus, hPGC-methylation of TEs is strongly associated with the subsequent acquisition of H3K9me3, as well as enhancer marks in human tissues. Notably, the initiation of the onset of lineage-specific histone marks therefore appears to begin in the germline with the retention of DNA methylation within hPGCs.

### TE-derived enhancers may facilitate the development of the human fetal brain

Since the enrichment of several TE subfamilies occurs within brain-specific H3K9me3-marked regions (see above, Fig. 2b, Extended Data Fig. 2a), we explored the role of hPGC-methylated TEs in the context of the human fetal brain. We re-analyzed data from the developing cortex^37^ and identified 29,679 syntenic regions, enriched for H3K27ac in both human and rhesus macaque, as putative active enhancers (Supplementary Table 6). The human brain has evolved markedly compared to the macaque, being approximately 4.8 times larger in size (adjusted for body weight)^38^, with a comparatively 35% larger neocortex^39^. We identified novel insertions of hPGC-methylated TEs in human within these orthologous enhancer regions. LTR12C was the most highly hPGC-methylated TE insertion found within human brain enhancers (Fig. 4a). Pathway analysis revealed an enrichment of LTR12C within genes controlling cell adhesion, neurogenesis, and astrocyte differentiation that are all associated with corticogenesis (Fig. 4c). Notably, LTR12C has been reported to be a primary source of novelty in primate gene regulation^19^, which can influence human corticogenesis. Accordingly, hPGC-methylation may be associated with the promotion of human specific enhancer formation within the developing brain and potentially other tissues.

**Fig. 4:**
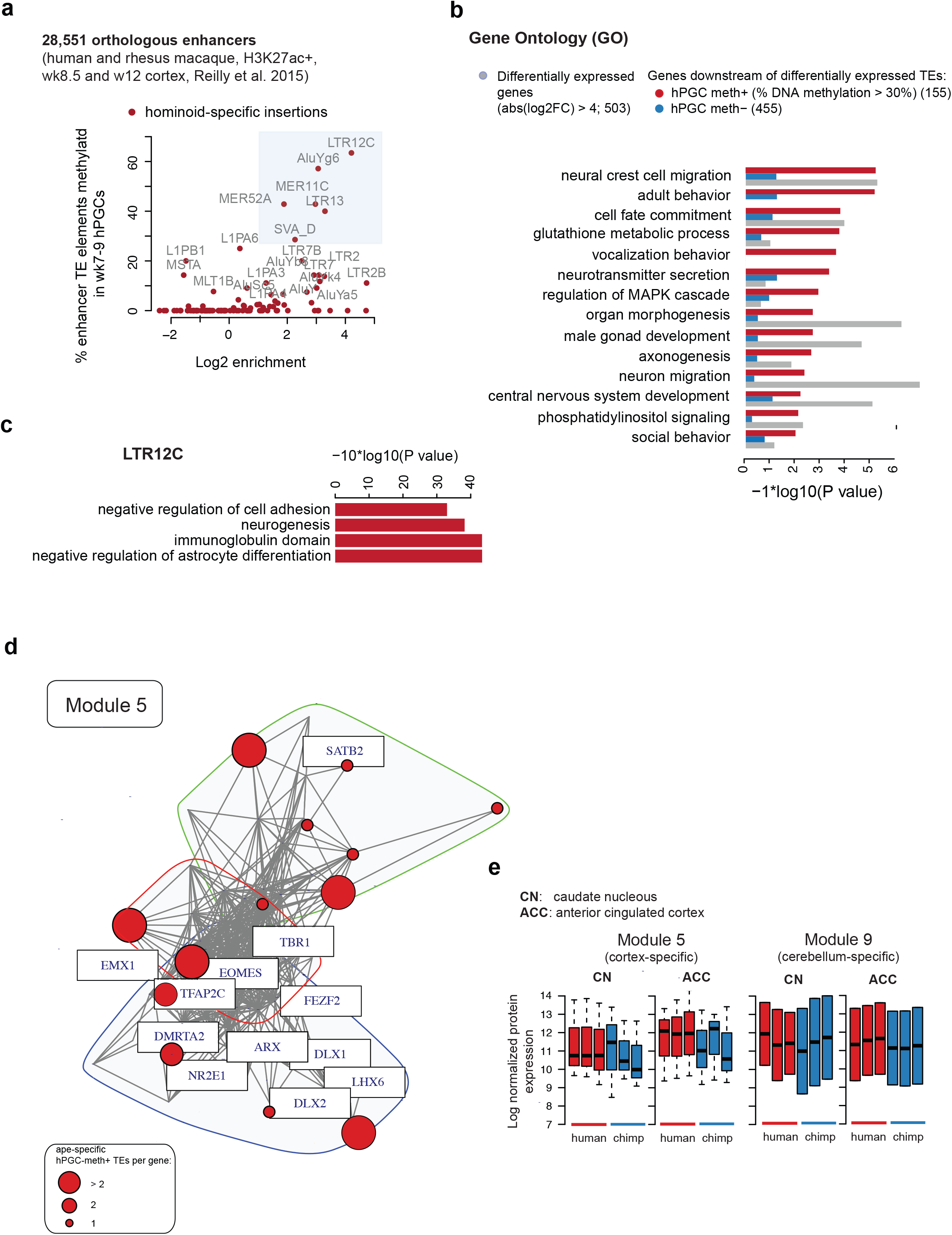
hPGC-methylated TE elements that switch to enhancers in the developing human brain. **a,** Scatter plot showing for each TE subfamily the percentage of elements that are DNA-methylated in wk7-9 hPGCs (y-axis) compared to the enrichment in 29,679 orthologous human and rhesus macaque enhancers (x-axis). Orthologous enhancers were defined by cross-referencing H3K27ac peaks determined at equivalent stages during corticogenesis (wk8.5 and wk12, data from Reilly et al., 2015, see Methods), with TE elements within the enhancers cross-referenced between human and rhesus macaque, respectively. **b,** Gene ontology (GO) term enrichment analysis for genes close to TEs that are differentially expressed between cortex and cerebellum (abs(log2FC) > 2; adj P value < 0.05). TEs that were within 50 kB distance upstream of the TSS or within the gene body of the genes were used for analysis. For each GO term red bars represent BH-corrected P values for genes that are close to hPGC-methylated TEs, blue bars for genes that are close to non-methylated TE elements, and grey bars represent P values for differentially expressed genes without considering TE information. **c,** Gene ontology (GO) term and Uniprot keyword enrichment analyses for genes that are within 50 kB distance of hPGC-methylated and ape-specific LTR12C elements in active enhancers of human corticogenesis. **d,** Top co-expression module (module 5) that is enriched for ape-specific hPGC-methylated TE insertions. The genes representative of the GO terms found to be enriched for this module are labelled. Nodes are highlighted and scaled according to the number of TE insertions close to the gene during primate evolution. The module was partitioned into overlapping communities using the Louvain method (see Methods). **e,** Bar plots showing the evolutionary divergence of protein expression in human compared to chimp in the caudate nucleus (CN) and the anterior cingulate cortex (ACC) for two co-expression modules. The average expression of the genes in each module is shown for three replicates per species (data from Bauernfeind et al., 2015).

For further support, we examined the relationship between TEs and the gene expression network of the fetal brain. We analysed 284 RNA-seq datasets from differing brain regions in the cortex and cerebellum of 172 fetal brains (post-conceptual week 4-17; mean 10.05 weeks)^40^. Identifying 20 unique modules in 6,766 differentially expressed genes (log2FC > 1, P value < 0.05), we observed that ape–specific hPGC-methylated TEs were primarily enriched in cortically expressed modules compared to those in the cerebellum (Extended Data Fig. 4a,b, Supplementary Table 7); the latter is significantly less diverse, functionally and morphologically, in humans compared to the cortex ^41, 42^. Within module 5, the most preferentially cortically expressed module, the insertion of two or more hPGC-methylated TEs within multiple nodes that correlate with gene expression (Fig 4d). These genes include *EOMES* expression in intermediate radial glial progenitor cells as they enter a proneural state ^43^, and *EMX1* that is critical for neocortical patterning^44^. Together they are significant for the development of human cortical gyrification, which is an adaptive evolutionary process in mammals facilitating cognitive development^45, 46^. In another cluster, *DMRTA2* and *DLX2* are both associated with at least one hPGC-methylated TE, and these genes are involved in neuronal migration and the switch between neuronal and oligodendrocyte cell fate, respectively^47, 48^.

Together, these observations show that hPGC-methylated regions exist at sites associated with networks that shape components of the unique architecture of the human brain, and within enhancers of cortical development, and gene-expression modules active in human neurodevelopment.

### The potential impact of hPGC-methylated TEs in the human cortex

Genes within Module 5 were more highly expressed in the developing human cortex than in chimps (Fig. 4d,e), prompting us to investigate several facets of hPGC-methylated and non-methylated TE expression in this context. Comparing 284 RNA-seq datasets from differing brain regions in the cortex, cerebellum and midbrain of 172 fetal brains (275 samples, see above), we observed 10,346 expressed TE elements (irrespective of DNA methylation status in PGCs), which exhibited clear spatiotemporal dynamic expression (Extended Data Fig. 4c, PC2 vs. PC1 and PC4 vs PC3). Young hPGC-methylated TEs (those shared with great apes or ‘ape-specific TEs’) such as L1PA2 (~ 7.6 million years old) showed high cortical expression, compared to L1PA4-11 families (~ 13 to 53 million years old^29^). Notably, the expression of the latter was preferentially observed within the cerebellum (Extended Data Fig. 4d). When comparing genes that are in close proximity (<50kb) to all differentially expressed TEs (log2FC > 2), we observed that hPGC-methylated regions were strongly associated with critical developmental processes in comparison to non-methylated TEs (Fig. 4b). One notable example is provided by *FOXP2^49^*, which has facilitated language development during human evolution^49,50^. Accordingly, these data further demonstrate that hPGC-methylated regions are strongly enriched within enhancers and gene-networks that are primarily active during corticogenesis, and importantly with those that show differential expression within the developing cortex.

### KRAB zinc finger proteins may mediate the exaptation of TEs as developmental enhancers

Next, we determined potential mediators that influence the enhancer activity of hPGC-methylated TEs in the brain. In particular, we considered the role of KZFPs, which binding to TEs to repress their activity in human embryonic stem cells (ESCs) and the early embryo^16,51^. Through their ability to recruit KRAB-associated protein 1 (KAP1) complex, histone-deacetylase-containing NuRD complex, histone-lysine methyltransferase SETDB1 and DNA methyltransferases ^29^, they also mediate methylation and chromatin modifying effectors^29^ (Extended Data Fig. 5a).

Analysis of genome-wide binding to TEs (ChIP-exo) for 236 KZFPs^7^ showed that multiple KZFPs, including ZNF93 and ZNF534, retain preferential binding for hominoid-specfic methylated TEs (L1HS and L1PA2, and HERVH, Extended Data Fig. 5b). When considering the binding of KZFPs on all TEs (methylated and non-methylated) by age, we observed a clear propensity for KZFPs to bind to the majority of young TEs (L1HS – L1PA3) irrespective of their methylation status, before subsequent continued enrichment of KZNFs binding to hPGC-methylated TEs (Extended Data Fig. 5c, Supplementary Table 8). It is possible that KZFPs may initially function primarily to suppress TE activity, but over time, KZFP-TE interactions may function to provide a layer of DNA methylation and possibly chromatin modifications at KZFP-TE sites. Although further experimental is required, it is possible that if such epigenetic modifications could then be transmitted to the totipotent zygote at fertilization, then this may offer a potential mechanism for intergenerational epigenetic inheritance.

After establishing KZFPs that are expressed in week 7-9 hPGCs and dynamically expressed in the brain, we found a subset of KZFPs with high expression within the cortex, including ZNF534, ZNF257 and ZNF98 (Fig. 5a). Moreover, we detected multiple examples of KZFP-TE interactions, where the expression of the KZFP in the cortex and cerebellum was significantly anti-correlated with that of the hPGC-methylated TEs, to which they have binding motifs, in particular for members of the LTR12 and SVA families (Fig. 5b, Supplementary Table 9). Some of these dynamically expressed hPGC-methylated TEs provides further evidence of KAP1 binding, and binding of the corresponding KZFP by ChIP-exo (Fig. 5c), We also found associated correlated expression of overlapping or near-by genes, which, as above, are enriched for those influencing neurodevelopmental processes; and this correlation was particularily strong for hPGC-methylated TEs with evidence for KZFP binding (Extended Data Fig. 5d). Thus, we propose that dynamically expressed KZFPs within the brain may bind to their corresponding TE motifs and influence localized gene expression, generating a somatically-acting exapted KZFP-TE network within the brain as observed in other cell types^16^. Crucially however, the onset of this process may begin within hPGCs and not entirely somatically; accordingly, the nidus for these networks may be inherited from the germline.

**Fig. 5:**
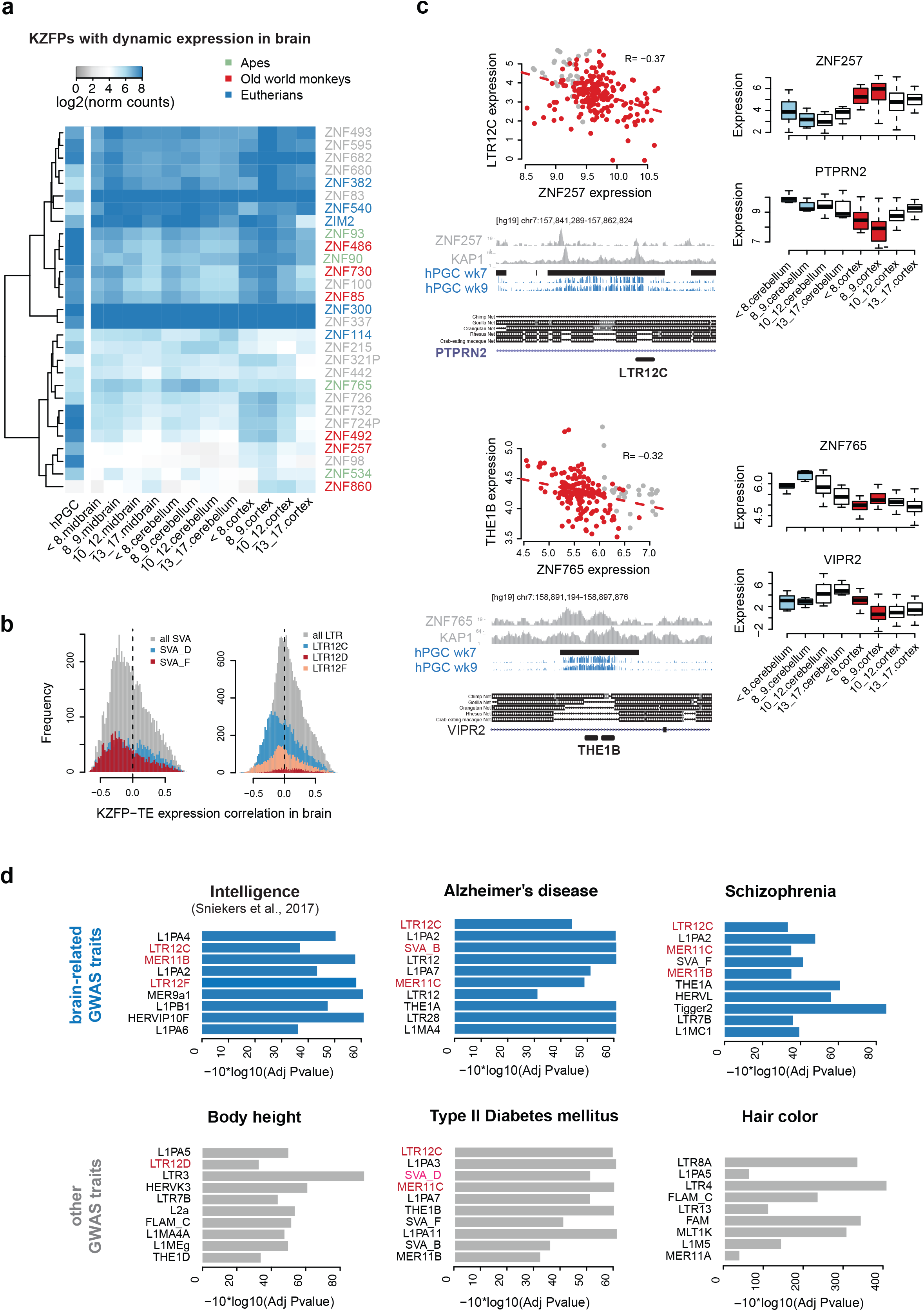
KRAB-ZFPs may influence the enhancer activity of hPGC-methylated TE elements in the brain. **a,** A heat map showing the average expression of KRAB zinc finger proteins (KZFPs) in wk7-9 hPGCs and in the developing brain. 29 KZFPs are differentially expressed between the cortex and cerebellum at any developmental stage, or between different stages of cortical development (CS22-wk17, 219 samples; log2FC > 1; P value < 0.05). **b,** Histograms showing the Pearson correlation coefficients between the expression of KZFPs and the expression of hPGC-methylated TEs of selected LTR and SVA subfamilies in the cortex and cerebellum (CS22-wk17, 219 samples). Selected enhancer-related TE subfamilies (such as LTR12C) are highlighted in red; and their expression is predominantly anti-correlated to KZFP expression. **c,** Representative genome browser tracks showing for TE elements, (bottom left) the KZFP and KAP1 ChIP-seq coverage, the % DNA methylation in wk7-9 hPGCs, and the conservation in primate genomes. (top left) Scatterplots show the expression of the TE element versus the expression of the KZFP in CS22-wk17 cortex and cerebellar samples. TE expression and KZFP expression are anti-correlated. Boxplots show the average expression of the gene close to the TE element (bottom right) and the expression of the KZFP (top right) in CS22-wk17 cortex and cerebellar samples. **d,** Bar plots showing the gene-based binomial P values for the association between selected brain-related and other/metabolic traits and TE subfamilies (see Methods). TE subfamilies, where > 40% of TE elements are hPGC-methylated and > 70% of TE elements are only detected in apes, are shown in red. See also Extended Data Fig. 6a and Supplementary Table 10. TE subfamilies are ranked by the number of genes that close to the GWAS SNPs.

### Association of hPGC-methylated regions with human traits and diseases

Finally, we examined if some TE subfamilies may be associated with common human diseases. Comparing the overlap between methylated TEs and genes identified in GWAS loci associated with human diseases and traits, we identified 294 TE subfamilies that were enriched for at least one human trait or disease (Supplementary Table 10), including height and intelligence, and diseases including schizophrenia and Alzheimer’s disease (Fig. 5d). Particular TE classes were enriched for a heterogeneous array of traits or diseases affecting multiple organs, were families such as LTR12C enriched conditions including type 2 diabetes mellitus, oesophageal carcinoma and Alzheimer’s disease. In contrast, other families were associated primarily with single organ conditions such as LTRA8, which was enriched for dermatological traits and conditions such as hair colour, vitiligo and sensitivity to sun (Extended Data Fig. 6a). These observations suggest that hPGC-methylated regions may be associated with mediating both adaptive traits and also potentially several human diseases. Altogether evidence suggests that hPGC-methylated regions are likely to have an impact on the regulation of genes that have significant functions in human health and disease. Speculatively, variations in hPGC methylation within specific TE families may constitute a risk factor for certain disorders, which may partly explain the elusive nature of heritability of some conditions and traits^52^.

## DISCUSSION

Our study provides a foundation for testable hypotheses that the transgenerational inheritance of loci, which are resistant to the acute epigenetic-resetting in the human germline, might have a significant role in human evolution, development, and disease. It is critical to establish whether the persistent methylation of transposable elements within the human germline is a stochastic process, or an adaptive component of the human germline, which primarily involves hominoid-specific TEs^1^. The exact mechanism and mediators of resistance to DNA demethylation in the acutely hypomethylated hPGCs remain to be determined, but KZFPs binding is amongst a possibility to mediate this process^53^. KZFPs binding to young TEs appears prevalent regardless of their DNA methylation status, as well as to older TEs that are no longer mobile. The apparent progressive increase in the proportion of TEs with methylation in various subfamilies is noteworthy. The basis for such an outcome merits consideration, together with the impact of the persistent TE-methylation and their transmission, and their implications for evolutionary biology, human development and disease.

The TE family-specific methylation reprogramming within gametes and the significant methylation in hPGCs correlate with reprogramming of H3K9me3. Their enrichment with heterochromatin marks in early embryonic stem cells and their *in vivo* counterpart may provide the nidus for the onset of somatic epigenetic programming during human development^54^. Notably, the timing of these events could coincide with the rapid peri-implantation epiblast proliferation and selection through cell competition, before the onset of somatic and germ cell fates. For example, the marked association of methylated ERVs and LTRs (such as LTR12C or MER52A) with H3K9me3 erasure in primed ESCs, and whether KZFP binding at this point plays a role, remains to be established. Conversely, hPGC-methylated SVA elements, including very young human-specific elements such as SVA_E and SVA_F, which lack tissue-specific histone marks, show a strong increase of H3K9me3 in primed pluripotent cells. Such behaviour in the early phase of development, where cells gain a competent state for all three germ layers and the germline, may ensure the co-association with H3K9me3 in a relatively homogeneous fashion; subsequent selection processes may act to retain H3K9me3 marks in a tissue specific manner.

A recent study suggested that Krüppel-like transcription factors, including KLF4 can activate the embryonic genome in early embryos through binding to young TEs and inducing their expression; subsequent expression of KZFPs then represses the TEs^16^. Similar events occur in naïve hESCs that contain relatively hypomethylated DNA compared to primed hESCs. Since at least some KLF family members are expressed in early human PGCs undergoing DNA demethylation^55,56^, this might suggest that a similar feedback mechanism could operate in the human germline. In addition, the most recently acquired ERV (HERVK) expression is also detected and driven by OCT4 in human preimplantation embryos that ceases after implantation^57^. The hominoid-specific young TEs in the germline, and their exaptation may provide species-specific regulators of gene expression in multiple tissues.

We provide a potential basis for somatic modulation of hPGC-methylated TEs within the brain. Accordingly, young methylated TEs have become enriched within a variety of modules that can apparently influence neurodevelopmental processes, in particular cortical developmental networks. These networks incorporate both enhancers (such as LTR12C or SVAs) and TEs associated with repressive histone complexes such as H3K9me3. Within 5 million years of their incorporation into the human genome (based on genome age), we detect exapted somatic TE networks, which merits further investigation. With more precise information on TE loci within the genome, high quality cell-specific brain RNA-seq datasets could reveal their influence on human brain development as well as in other somatic tissues. An association between methylated TEs in almost 300 families and several human traits and diseases suggests that these TEs may influence a multitude of adaptive human traits across multiple organ systems and tissues together with several human diseases.

It remains a possibility that the hominoid-specific TE loci that are resistant to reprogramming and retain methylation in hPGCs, constitute a layer of heritable information, with an impact on human evolution, development and disease.

## Acknowledgements

We thank members of the Surani lab for helpful discussions, as well as Drs Felipe Karam Teixeira, Thorold Theunissen, and Jose C. Silva for feedback on this work. SD received support through a core grant to the Wellcome Trust – Medical Research Council Stem Cell Institute, and through a grant to AS, MJK is an NIHR-funded Academic Clinical Lecturer, WWT receives support by a Croucher Foundation Fellowship and by the Newton Trust, EM from the Icelandic Research Fund (174564-053), PFC is a Wellcome Trust Principal Research Fellow (212219/Z/18/Z), and a UK NIHR Senior Investigator, who receives support from the Medical Research Council Mitochondrial Biology Unit (MC_UU_00015/9), the Medical Research Council (MRC) International Centre for Genomic Medicine in Neuromuscular Disease, the Evelyn Trust, and the National Institute for Health Research (NIHR) Biomedical Research Centre based at Cambridge University Hospitals NHS Foundation Trust and the University of Cambridge. The views expressed are those of the author(s) and not necessarily those of the NHS, the NIHR or the Department of Health. AS is a Wellcome Senior Investigator with a grant from the MRC. The Gurdon Institute receives a core grant from the Wellcome Trust and Cancer Research UK.

## Author contributions

SD and AS conceived the project. SD designed and performed bioinformatics analyses. MJK, WWT and EM analysed the data and contributed to the discussions of the results, and TK and PFC provided essential resources. SD, MJK, and AS wrote the manuscript with input from all authors.

## Extended Data

**Fig. 1: Germline expression and age of hPGC-methylated TEs**. **a,** A pie chart showing the intersection of hyper-methylated regions determined from two independent studies of DNA methylation in early hPGCs (see Methods, data from Tang et al., 2015 and Guo et al., 2015). **b,** A scatter plot showing for each TE subfamily the percentage of elements that are expressed in wk7-9 hPGCs (log2(normalized read counts) > 5), and the percentage of elements that are methylated in wk7-9 hPGCs (DNA methylation > 30%). Alu and LINE L1 elements are shown in a subfigure for clarity. Names are shown for TE families, of which at least 5% of annotated elements are methylated in wk7-9 hPGCs. **c,** A scatter plot showing for each TE subfamily the percentage of elements, which can be mapped to a syntenic region in chimp, gorilla, orangutan or gibbon genomes, and the percentage of elements that are methylated in wk7-9 hPGCs (DNA methylation > 30%). Predominantly hominoid-specific TE subfamilies, which retain DNA methylation in hPGCs, are highlighted in blue.

**Fig. 2: Association between hPGC-methylated regions and tissue-specific histone marks. a,** A heat map showing the enrichment (Z-scores) for tissue-specific H3K9me3 ChIP-seq peaks (data from NIH Roadmap Epigenomics Project) in 9,902 human-specific hPGC-methylated TE elements that cannot be mapped to syntenic regions in non-human primates or other mammals, compared to 1,000 random sets. Only TE families with a Z-score of at least 10 for one cell type are shown. **b,** Related to Fig. 2c. Bar plots showing for each TE subfamily the percentage of elements that intersect with H3K9me3 and enhancer-related (H3K4me1+H3K27ac) ChIP-seq peaks in 7 fetal and 28 adult tissues (data from NIH Roadmap Epigenomics Project) for TE elements that are methylated (> 30 % DNA methylation) or unmethylated in wk7-9 hPGCs. TE subfamilies with significant differences in enhancer-related (H3K4me1+H3K27ac) peaks (P value < 1e-5, Pearson’s Chi-squared test) for hPGC-methylated vs non-methylated regions are coloured in red. Left: bar plot showing the fraction of TE elements for each subfamily that are methylated in wk7-9 hPGCs (DNA methylation > 30%). TE subfamilies are ranked according to the fraction of hPGC-methylated elements per TE subfamily.

**Fig. 3: DNA methylation of hPGC-methylated TE elements in gametes. a,** Bar plots showing for selected TE subfamilies the percentage of hPGC-methylated elements that are not methylated (< 20 % 5mC), partially methylated (>= 20% and < 80%), or methylated (>= 80%) in sperm, oocytes or blastocysts (data from Okae et al., 2014). TE subfamilies are ranked by the 5mC variability (coefficient of variation) in sperm (see also Supplementary Table 5). Older TE subfamilies are printed in bold. **b,** Density scatter plots of % DNA methylation in sperm compared to oocytes, and in blastocysts compared to wk7 hPGCs for TE elements that are methylated (DNA methylation > 30%) in wk7-9 hPGCs. **c,** Heatmaps showing the % DNA methylation in wk7-9 hPGCs, oocyte, sperm and blastocysts (data from Okae et al., 2014) for TE elements that are methylated (DNA methylation > 30 %) or unmethylated in wk7-9 hPGCs. **d,** A scatter plot showing for each TE subfamily the average H3K9me3 logFC in primed vs naïve hESCs (data from Theunissen et al. 2016) for TEs that are methylated in hPGCs (x-axis) compared to TEs that are non-methylated (y-axis). For instance, for the LTR12C subfamily in the topleft quadrant, hPGC-methylated TE elements lose H3K9me3 on average, but non-methylated TE elements of the LTR12C gain H3K9me3 in primed hESCs.

**Fig. 4: Co-expression module analysis in the human fetal brain. a,** A co-expression module analysis of 6,766 genes differentially expressed between cortex and cerebellum at the same developmental stage, or between different developmental stages of cortical development (219 samples, abs(log2FC) > 1, adj P value < 0.05) using k-means clustering of the first 25 principal components (see Methods and Supplementary Table 7): (Top) A bar plot showing the number of genes in each of the 20 kmeans-derived modules. (Bottom) A PCA plot depicting the average PC scores of the genes in each co-expression module. Modules are coloured by their average log2FC between wk8-9 cortical and cerebellar samples with red modules being up-regulated in the cortex, and blue modules being up-regulated in the cerebellum. **b,** A bar plot showing the enrichment of hPGC-methylated TE elements close to the genes in each kmeans-derived co-expression module (see Methods). TE elements are classified according to the primate lineage in which they first emerge. Modules are ranked by their enrichment of ape-specific TE elements. **c,** A principal component analysis (PCA) of TE expression during cortex, midbrain and cerebellar development (275 samples, CS14 to wk17). TE elements with at least log2(normalized read counts) > 5 were included in the analysis. Samples are coloured according to their eventual brain region in the adult brain and their developmental stages at the time of RNA extraction. hPGC wk5.5-wk7 samples are shown in black. **d,** Heat maps showing the expression of TE elements that are differentially expressed between cortex and cerebellum, or between different developmental stages of cortical development (abs(log2FC) > 2, adj P value < 0.05). In total, 9,363 TE elements were differentially expressed.

**Fig. 5: Correlation between the expression of hominoid-specific KZFPs and TEs in the developing human brain**. **a,** A schematic showing that KAP1 can be recruited to specific sites by KRAB zinc finger proteins. KAP1 acts as a scaffold for the assembly of a large repression complex consisting of the NuRD histone deacetylase complex and the H3 lysine 9-specific histone methyltransferase SETDB1. KAP1 further interacts with HP1, which in turn can bind to histone H3 that has been methylated on lysine 9 (H3K9me3). **b,** Ape-specific KZFPs that are expressed during human embryonic brain development. (Left panel) A heat map showing the enrichment (Z-scores) of KZFP ChIP-exo peaks (data from Imbeault et al., 2017) in 9,902 human- and 52,654 ape-specific hPGC-methylated TE elements, compared to 1,000 random sets. Only those KZFPs and TE subfamilies that have a Z-score of at least 10 are shown. KZFPs that are differentially expressed between cerebellum and cortex at any developmental stage, or between different stages of cortical development are shown in blue (see also Fig. 5a). (Right panel) % sequence identity of the human KZFP gene to its one-to-one ortholog in non-human primates. **c,** Bar plots showing the percentage of TEs in each subfamily that intersect with KZFP ChIP-exo peaks (data from Imbeault et al., 2017). TE elements are categorized as methylated (DNA methylation > 30%) or non-methylated in wk7-9 hPGCs. KZFP binding is more prevalent for hPGC-methylated in comparison to non-methylated elements of the same TE subfamily (P value < 0.01, Pearson’s Chi-squared test), indicating that KZFPs might regulate the protection of hPGC-methylated TE elements from DNA de-methylation. Left: bar plot showing the fraction of TE elements for each subfamily that are methylated in wk7-9 hPGCs. **d,** Bar plots showing the percentage of TEs that have a significant correlation in expression (Pearson correlation coefficient > 0.5, P value < 0.001) with their neighbouring genes (distance < 15 kB) for hPGC-methylated and non-methylated TEs. TE elements are further categorized into three classes: (1) those intersecting with H3K9me3 ChIP-seq peaks, (2) those intersecting with enhancer-related (H3K4me1+H3K27ac) ChIP-seq peaks, and (3) those intersecting with enhancer-related ChIP-seq peaks and with at least two KZFP peaks (data from NIH Roadmap Epigenomics Project, and from Imbeault et al., 2017).

**Fig. 6: Association of hPGC-methylated TEs with human traits and diseases. a,** Bar plots showing for selected hominoid-specific TE subfamilies the gene-based binomial P values for the association with the top 10 most significantly associated GWAS traits. GWAS traits are ranked by the number of genes shared with the TE subfamily.

## Methods

### DNA methylation analyses

#### Description of data sets

DNA methylation data sets were obtained from public sequence archives for week 10-11 hPGCs (Guo et al., 2015; GEO: Accession: GSE63818), for gametes (oocyte and sperm) and blastocysts (Okae et al., 2014; Japanese Genotype-phenotype Archive: Accession: JGAS00000000006), and naïve and primed hESCs (Guo et al., 2017; GEO Accession: GSE90168). Our previously published data set from wk7-9 hPGCs (Tang et al., 2015) and data set from wk10-11 hPGCs from Guo et al. (2015) both represent the most de-methylated stage in primordial germ cells. Naïve hESCs most closely resemble the blastocyst stage, and both data sets represent a further wave of DNA de-methylation in the pre-implantation embryo.

#### Data processing

PBAT reads were pre-processed by removing 4nt from the 5’ end of the first read, and 15 nt from the second read to remove random primers using *Trim Galore!* (parameters: ‘--paired --clip_R1 4 --clip_R2 15 -three_prime_clip_R1 1 - three_prime_clip_R2 1 -q 30’). Quality-trimmed reads were aligned to the human reference genome (GRCh37/hg19 and GRCh38/hg38) using *Bismark* (parameters: ‘-n 2 -l 100 --bowtie1 –pbat’). Duplicates were removed using *Bismark deduplicate*. Alignments were further processed with *MethPipe*. Bam files were converted by *to-mr*, and methylation levels on CpGs were determined with *methcounts*, and symmetric CpG sites on opposite strand were merged by *symmetric-cpgs*. UCSC genome browser tracks (bigwig files) were generated by calculating a smoothed average of the DNA methylation levels of CpGs on 100nt-sliding windows (step 10 nt) using custom Perl scripts. Methylation levels of the hypo-methylated hPGC genomes were inverted (1- % methylation), and hyper-methylated regions were determined using the *Methpipe hmr* command. Hyper-methylated regions obtained from individual and pooled replicates were merged, and only hyper-methylated with at least 30% methylation in one individual or pooled replicate were retained. Published DNA methylation data sets were processed in the same way, except that end repair was not performed on WGBS-seq reads, and they were aligned with the ‘–directional’ option.

### ChiP-seq analyses

ChiP-seq data sets for KAP1 and H3K9me3 were obtained from the NCBI Gene Expression Omnibus (GEO) for human naïve and primed hESCs (Theunissen et al., 2015; Accession: GSE75868), and ChIP-exo data sets for 238 KZNFs in HEK 293T cells (Imbault et al., 2017; Accession: GSE78099). Reads were pre-processed with *Trim Galore!* (parameters: ‘--length 25 -q 30’), and aligned to the human reference genome (GRCh37/hg19 and GRCh38/hg38) using *bowtie* (parameters: ‘-M 1 -v2 –best --strata’ to filter unique alignments only. Narrow peaks were called with *macs2 callpeaks* (parameters: ‘-g 3e9 -q 0.05’).

### Analyses of transposable elements

#### DNA methylation

*RepeatMasker*-annotated TE elements were obtained for the GRCh37/hg19 and GRCh38/hg38 human genomes from the *UCSC Genome Table Browser*. % DNA methylation levels on transposable elements were determined by using *MethPipe roimethstats* considering only CpGs with at least 5X coverage. hPGC-methylated TE elements were required to have at least 30% methylation in one individual or pooled replicate, and to intersect with a hyper-methylated region by 50% of the length of the TE element.

#### Mapping of TE elements to non-human primate genomes

Pairwise alignments files with non-human primate genomes (and other mammals) were obtained *from UCSC Genome Browser Downloads*. Human *RepeatMasker*-annotated TE elements were mapped to the genomes of other species using *liftover*. TE elements which mapped by 0.9-1.1 of their length in human to the other species were considered to be conserved.

#### TE expression analysis in hPGCs and brain

RNA-seq reads were preprocessed using *Trim Galore!*, and reads from hPGC and brain RNA-seq data sets were trimmed to uniform length (50 nt) to be comparable. Reads were aligned to the human reference genome (GRCh37/hg19 and GRCh38/hg38) using *bowtie* (parameters: “-m 1 -v 2 –best --strata”) to filter the best and unique alignments only, in order to identify precise locations of TE expression. Read counts on TE elements were obtained by *featureCounts*, and differential expression analyses were performed using the R *Bioconductor DESeq* package. A principal component analysis for TEs with an expression of at least log2 (normalized counts) > 2 was performed using the R *prcomp* function (see below for further details on the brain RNA-seq data set).

#### Intersection with tissue-specific histone marks

Narrow peaks for 36 *in vivo* cell types, and for H1/H9 hESCs were obtained from the NIH Roadmap Epigenomics Project. The *in vivo* cell types were selected, for which enhancer-related histone marks (H3K4me3+H3K27ac), H3K9me3 and H3K36me3 were all available. For the H3K9me3 enrichment analysis in Fig. 2b, 1,000 random peak sets of the same size as the peaks called for the Roadmap Epigenomics data sets and within the vicinity of genes (< 50 kB) were generated with *bedtools shuffle* command. P values and Z scores for the enrichment of H3K9me3 were estimated by counting the overlap N with the 1,000 randomly generated peak sets, where the P value is defined as (N+1)/(1000+1). Pearson chi-squared tests in Fig. 2a (global) and 2c (TE subfamily-based), of observed proportions of TEs that intersect with histone marks for methylated vs non-methylated TEs were performed with the *R chisq.test* function.

#### Enrichment of hPGC-methylated TE elements in brain enhancers

ChIP-seq data sets for enhancer-related histone marks (H3K4me2 and H3K27ac) from the whole cortex at wk8.5 and primitive frontal and occipital tissues at wk12 for human and rhesus macaque were obtained from GEO (Reilly et al., 2015; Accession: GSE63649), pre-processed with the standard ChIP-seq protocol (see above), and aligned to the human (GRCh37/hg19) and rhesus macaque (rheMac8) reference genomes. Narrow peaks were called with *macs2 callpeaks*. Active enhancers in human and rhesus macaque were defined as H3K27ac peaks that intersected with an H3K4me2 peak, and did not intersect with an annotated promoter region. Human enhancer-related H3K27ac+H3K4me2 peaks were then mapped to the rhesus macaque genome by *liftover*, and human enhancers that intersected with an H3K27ac+H3K4me2 peak in rhesus macaque by at least 30% were considered conserved. We then cross-referenced and identified the transposable elements in the human enhancer regions that were not present in the orthologous region in macaque, and their DNA methylation in wk7-9 hPGCs, in order to identify hPGC-methylated TEs that were selected or newly inserted during human evolution and gain enhancer activity at this stage in human cortical development.

### Brain RNA-seq data analysis

#### Description of data sets

We analysed in total 275 cortex, midbrain, and cerebellum RNA-seq samples from the developing human brain by access to the HDBR resource^40^. The brain RNA-seq data set is unique in its age range (CS14-wk17), and number of brains (172) studied, in particular the samples earlier than wk8 of brain development. The majority of samples are from the time between wk9-12, where the major brain regions are established and the early stages of cortex development occur.

#### Data processing

Paired-end brain RNA-seq reads from the HDBR resource were trimmed of adapters using *Trim-Galore!.* Pre-processed reads were aligned to the human reference genome using *TopHat2*. Read counts on ENSEMBL protein coding, lincRNA and processed transcripts were obtained by *featureCounts*. Normalization and differential expression analyses were performed using the R Bioconductor DESeq2 package.

#### Gene co-expression modules in brain

We identified 6,766 genes differentially expressed between cerebellum and cortex, or between different stages of cortical development (CS14-wk17, 219 samples, normalized expression > 5 in one condition; abs(log2FC) > 1, adj P value < 0.05). First, we reduced the dimensionality of the gene expression matrix (genes with log2 normalized expression > 5 in at least 2 samples were selected) by performing a Principal Component Analysis (PCA) using the R *prcomp* function on the center-scaled gene expression values. We then performed a k-means clustering on the first 25 principal components of the 6,766 differentially expressed genes using the R *kmeans* function to define 20 gene co-expression modules. We calculated the average expression log2FC between wk8-9 cortex and cerebellum for the genes in each k means-derived module, since KZFPs show a distinct expression in the cortex at this developmental stage, and visualized the average log2FC of each module in PC2 vs PC1 space in Fig. 4a. Visualization of the adjacency matrix of a module (where nodes represent genes, and edges represent the Pearson correlation coefficient of gene expression between two genes in all cortex and cerebellar samples) in Fig. 4d were performed with the R *igraph* package. In addition, we performed the Louvain community detection method on each module using the *cluster_louvain* function of the *R igraph* package.

For each gene we determined the TE elements within 50 kB upstream of the transcriptional start site (TSS) or within the gene body. TE elements were classified according to their age depending on the non-human primate genome (human only, great apes or apes), where they were still conserved. The enrichment score for hPGC-methylated TE elements within gene co-expression modules according to their age in Extended Data Fig. 4b was determined as log2 (TE count by age / # of genes in module)/(TE count / # of all expressed genes).

### Analyses of KRAB zinc-finger proteins (KZFPs)

614 human genes with KRAB box (Pfam: PF01352) and C2H2-type zinc finger domains (Pfam: PF00096) and their gene orthology information for non-human primates, and mouse were obtained from ENSEMBL (version 96). Since hPGC-methylated TEs were predominantly hominoid-specific, we determined in Extended data Fig. 5b all KZFPs that have newly emerged or significantly evolved their gene structure in apes. We compared their mutual % sequence identity versus all non-human primate genomes and mouse. KZFPs with transcripts that share a mutual % seq id > 90% with any macacque transcript (including “many-to-many” orthologs) were excluded. We further required that KZFPs were differentially expressed between cerebellum and cortex, or between different stages of cortical development (CS14-wk17, 219 samples, normalized expression > 5 in one condition; abs(log2FC) > 1, adj P value < 0.05). For KZFP binding enrichment analysis in hominoid-specific TE subfamilies, 1,000 random regions of the same size as the TE elements were generated with *bedtools shuffle* command. P values and Z scores for the enrichment of KZFP peaks were estimated by counting the overlap N with the 1,000 randomly generated TEs, where the P value is defined as (N+1)/(1000+1). P values for the Pearson correlation coefficient between KZFP and TE expression in Fig. 5b were determined with the R *cor.test* function.

### Association of hPGC-methylated TEs and GWAS traits

The most recent GWAS catalog was obtained from EBI (https://www.ebi.ac.uk/gwas/docs/downloads, Jan 2019); additional data on loci influencing human intelligence were obtained from Sniekers at al., 2017. We considered only high-confidence GWAS SNPs with an adjusted P value < 1e-10. Direct overlap of hPGC-methylated TEs or hyper-methylated regions with SNPs were rare; we therefore identified the nearest genes of SNPs and hPGC-methylated TEs within 50 kB of their transcriptional start sites or within the gene body, and evaluated the statistical significance of the overlap between gene sets associated with GWAS traits and those associated with hPGC-methylated TEs for each TE subfamily using a binomial test. P values were determined by using the R *pbinom* function and adjusted for multiple testing using the Benjamini-Hochberg (BH) correction.

