## Supplemental figures for "Transposable elements resistant to epigenetic resetting in the human germline are epigenetic hotspots for development and disease"

#### Extended Data Figure 1

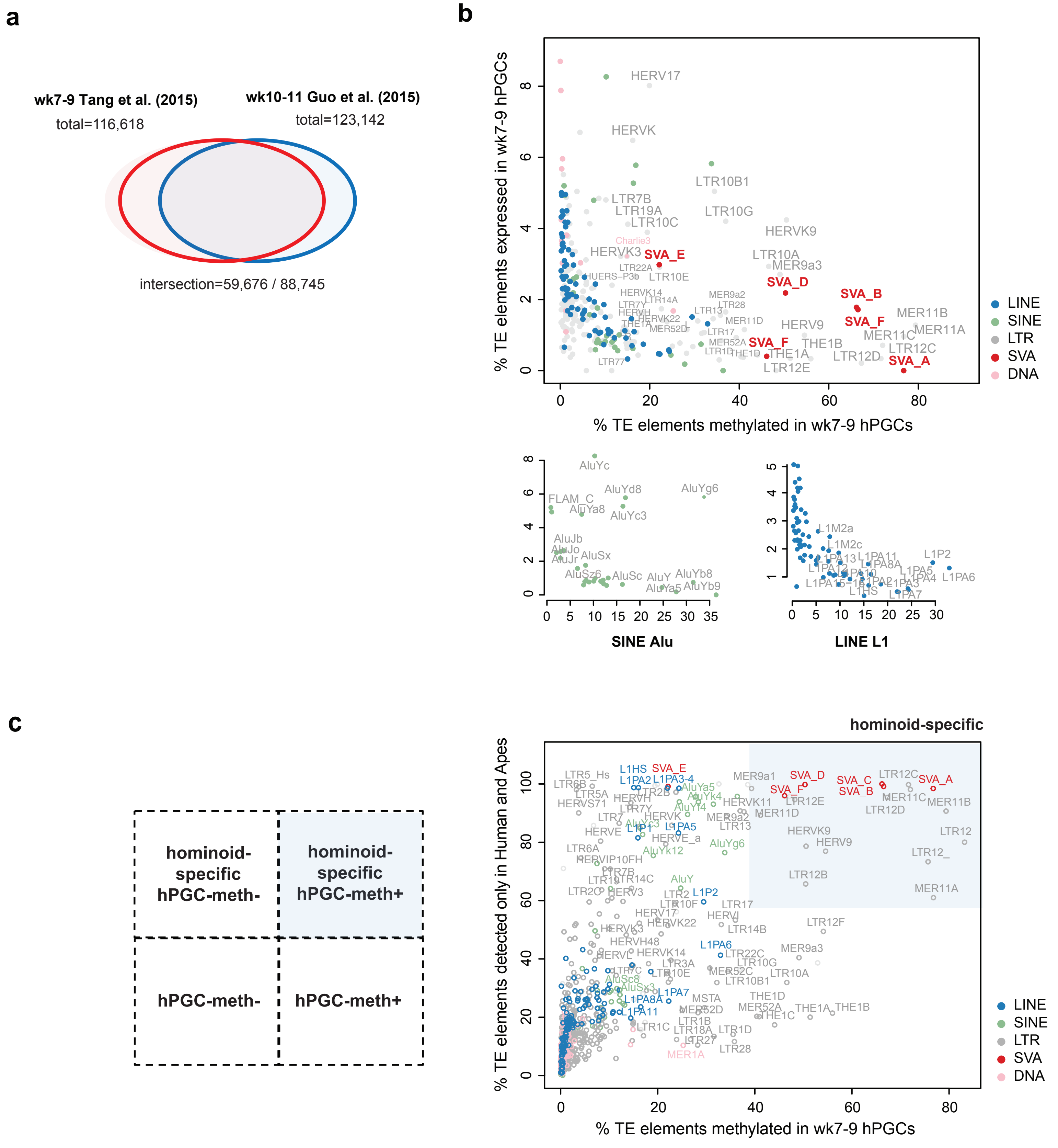

### Extended Data Figure 2

**a**

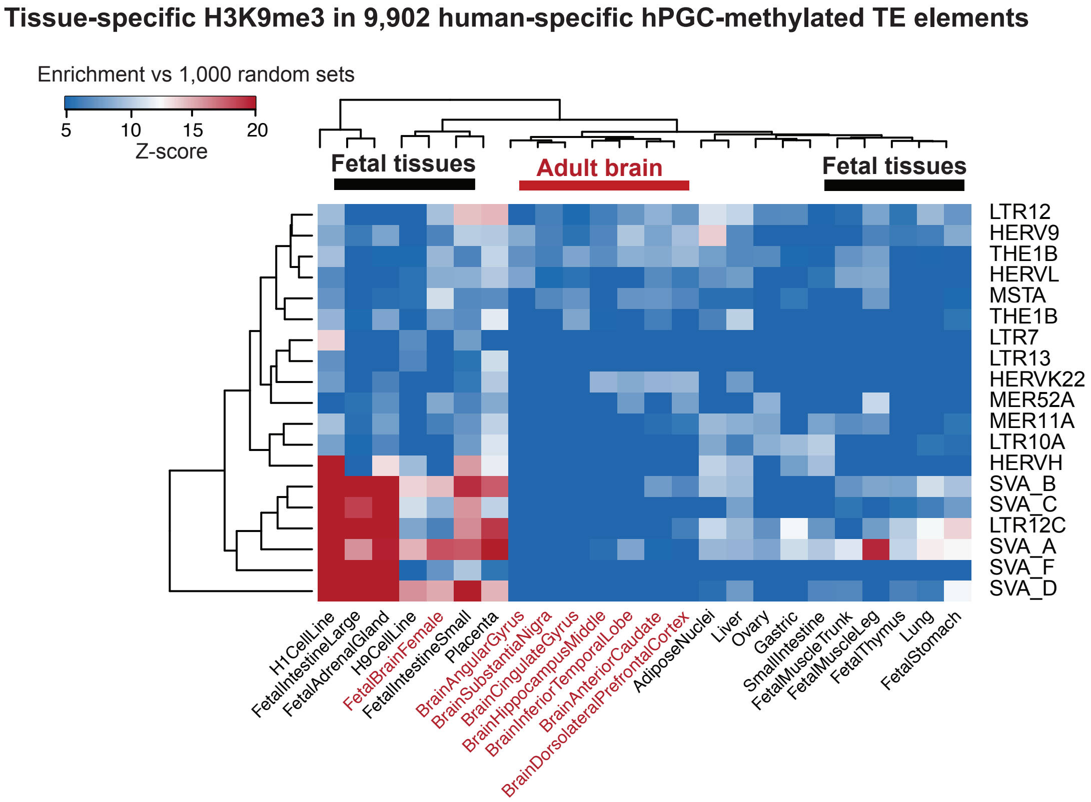


**b**

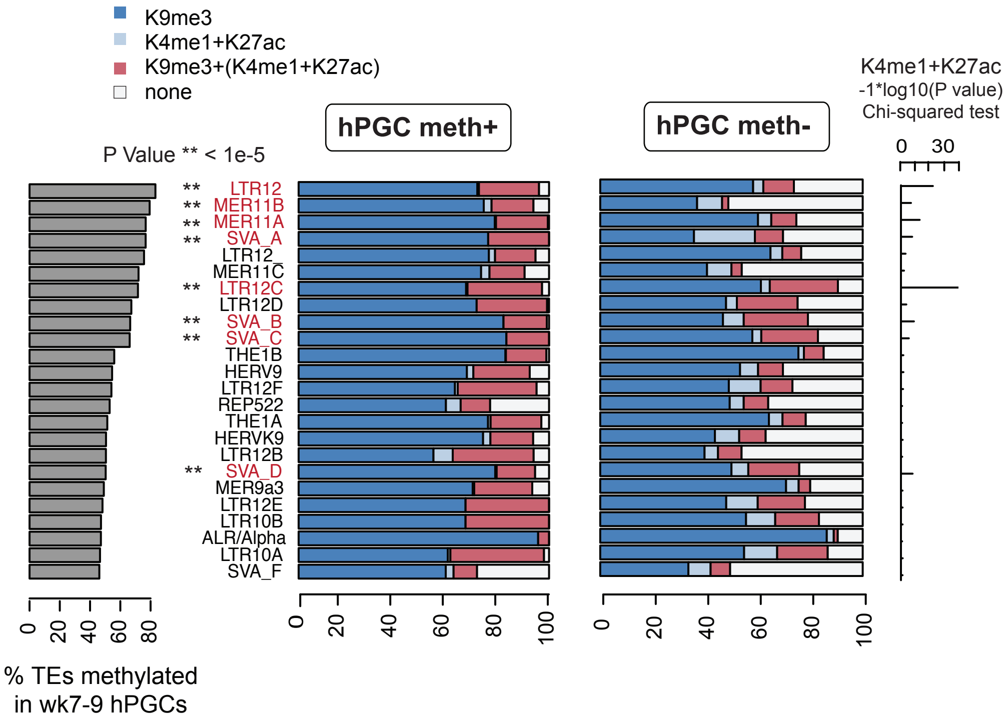

### Extended Data Figure 3

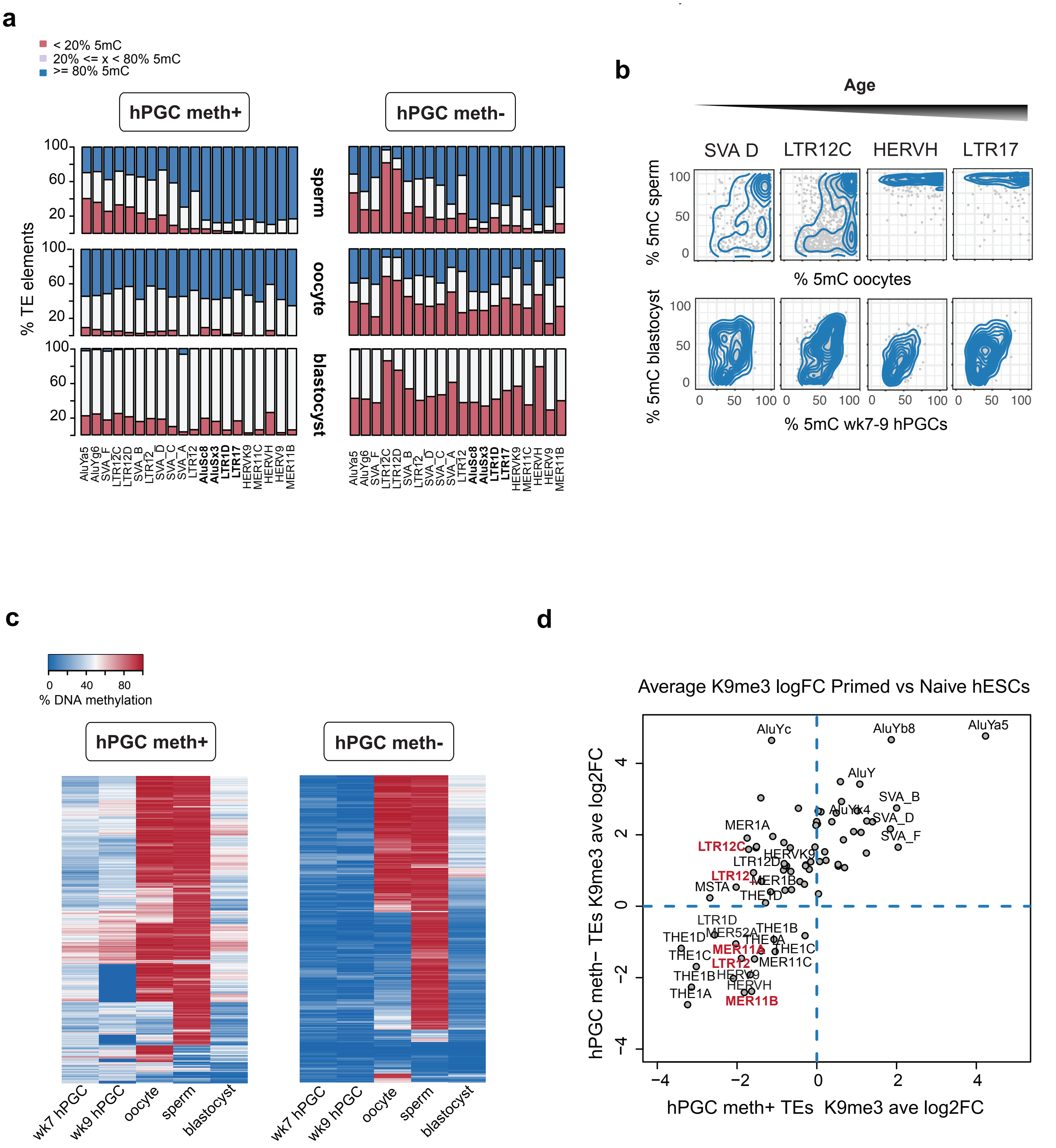

**Extended Data Figure 4**

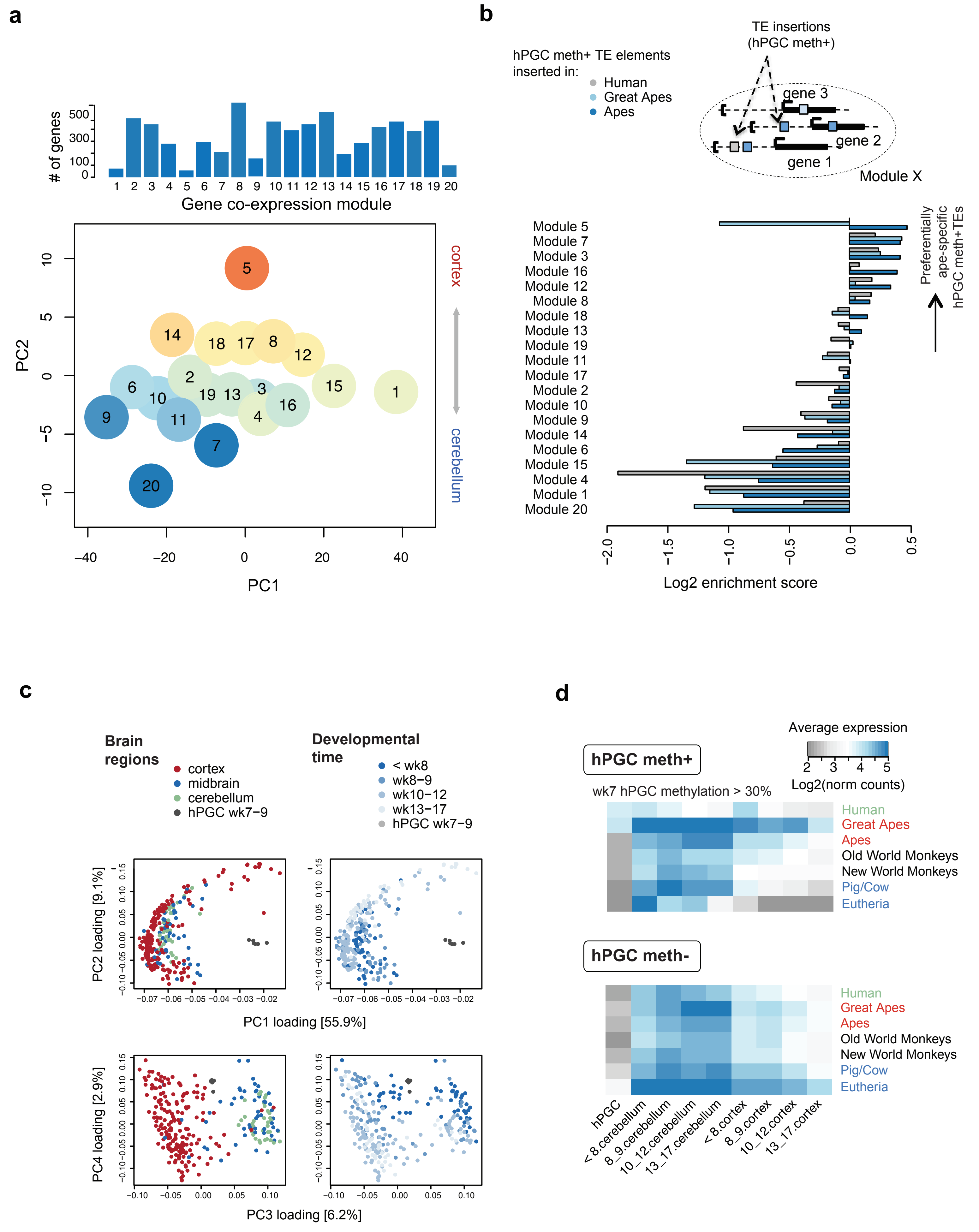

#### Extended Data Figure 5

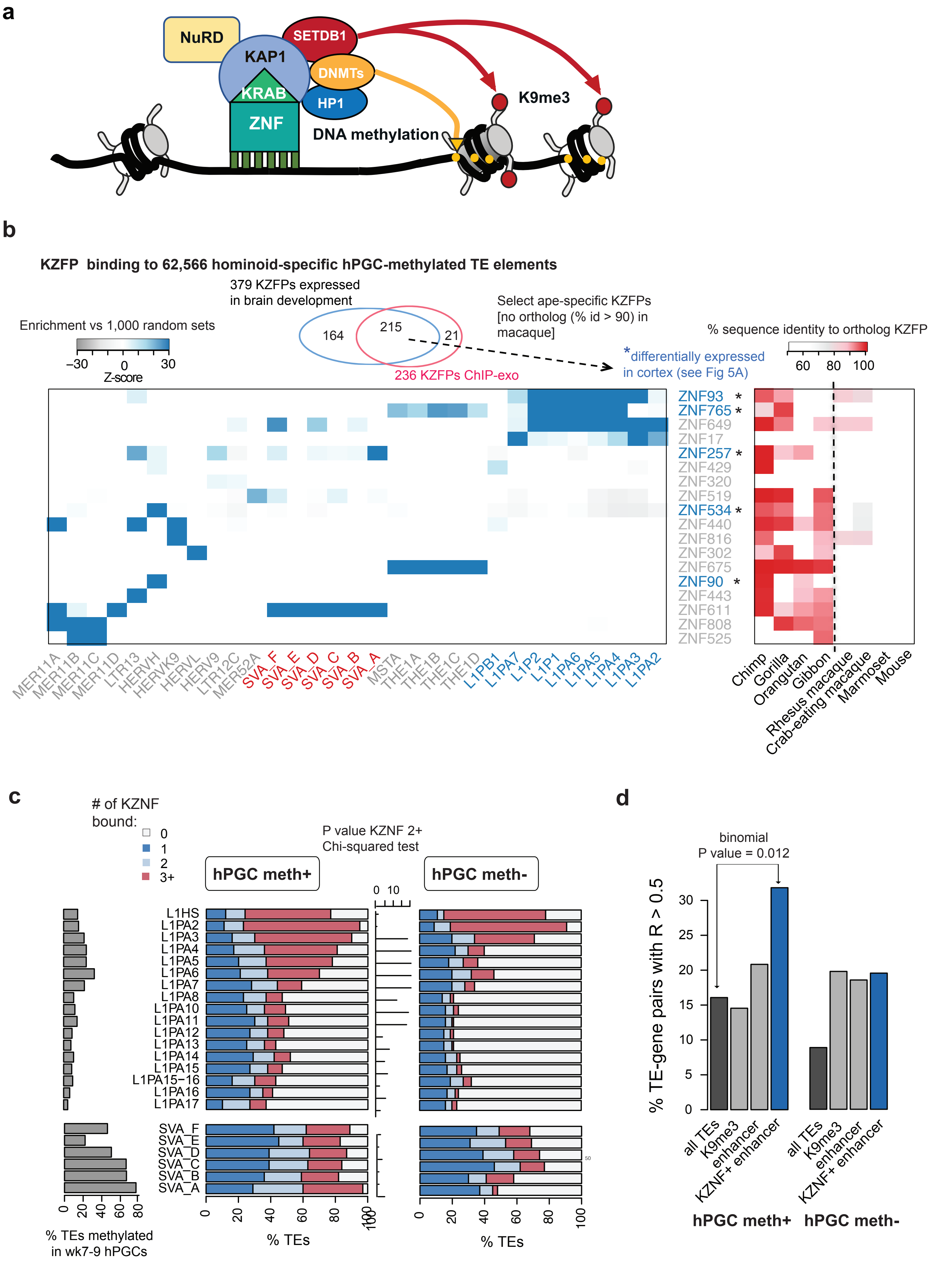

#### Extended Data Figure 6

**a**

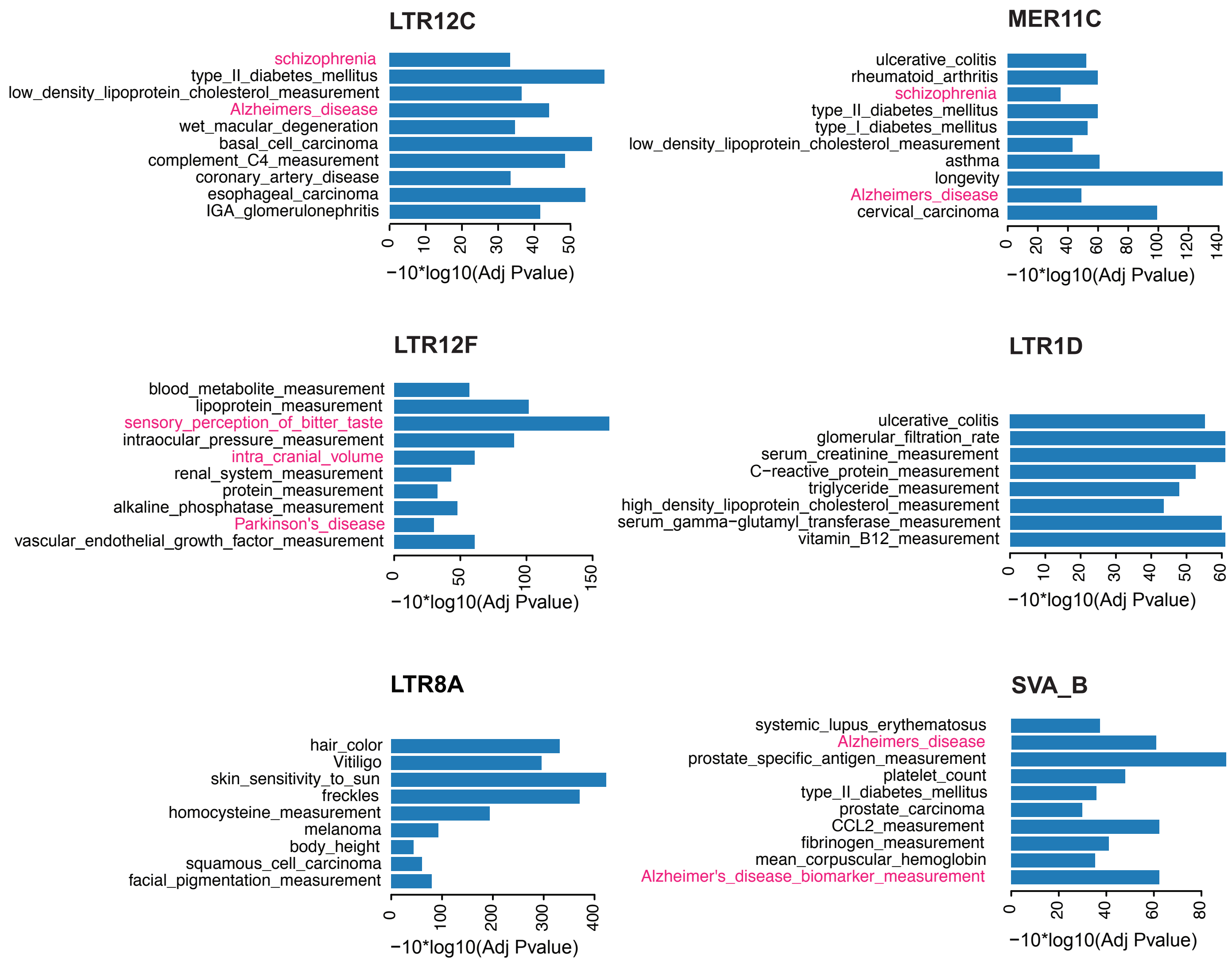
