## Supplemental Table Legends for "Transposable elements resistant to epigenetic resetting in the human germline are epigenetic hotspots for development and disease"

**Supplementary Tables**

**Table 1 Hyper-methylated regions in wk7-9 hPGCs.** DNA methylation in wk7-9 hPGCs; overlap with hyper-methylated regions from Guo et al. (2015). Data presented in Fig 1B, S1A.

**Table 2 TEs within wk7-9 hPGC hyper-methylated regions.** DNA methylation in wk7-9 hPGCs, gametes and blastocysts (data from Okae et al., 2014), and naïve and primed hESCs (data from Guo et al., 2017).

**Table 3 Conservation of TEs in non-human primates and other mammals.** TE lengths obtained in other genomes by liftover. Data presented in Fig 1C.

**Table** **4 Differences in tissue-specific H3K9me3 and enhancer-related histone marks (H3K4me1+H3K27ac)** **for hPGC-methylated vs non-methylated TEs**. Pearson’s Chi-squared test. Data presented in Fig. 2C, S2B.

**Table 5 Variability of DNA methylation for hPGC-methylated TEs in gametes and blastocyst.** Fraction of hPGC-methylated elements (DNA methylation > 30%) per TE subfamily, Coefficient of variation (CV; 5mC standard deviation/mean) in oocyte, sperm, and blastocyst for hPGC-methylated TEs. Data represented in Fig S3A.

**Table 6 Human cortical enhancer regions that are conserved in macaque.** 28,679 orthologous enhancer regions for the developing cortex (wk8.5-wk12) were identified by re-analyzing data from Reilly et al., 2015. Data presented in Fig. 4A.

**Table 7 Gene co-expression modules in the developing brain.** K-means cluster membership of 6,766 genes differentially expressed between the cortex and cerebellum (CS14-wk17, 219 samples, normalized expression > 5 in one condition; abs(log2FC) > 1, adj P value < 0.05). Data presented in Fig S4A.

**Table 8 Differences in KZFP binding for hPGC-methylated vs non-methylated TEs.** Pearson’s Chi-squared test. Data presented in Fig. S5C.

**Table 9 KZFP-TE expression correlation.** Correlation coefficient for the expression of dynamic KZFPs (see Fig. 5A) and hPGC-methylated SVAs and LTRs in the cerebellum and cortex (219 samples). Data presented in Fig. 5B/C.

**Table 10 Associations between hPGC-methylated TEs and GWAS traits.** Data presented in Fig. 5E and S6A.
